## Supplemental Figures for "Genetically-based sperm discrimination in the vaginal tract of a primate species"

Supplemental materials for **"Genetically-based sperm discrimination in the vaginal tract of a primate species"**

This supplement consists of:

**Supplementary Figures 1-7**

- Supplementary Figure 1. GSEA pathways enriched for differential expression in the fertile phase.
- Supplementary Figure 2. Vaginal pH does not vary significantly between cycle phases.
- Supplementary Figure 3. GSEA pathways enriched for differential expression post-copulation.
- Supplementary Figure 4. GSEA pathways enriched for differential expression in relation to MHC I allelic complementarity.
- Supplementary Figure 5 Leave-one-out sensitivity analysis of vaginal pH model estimates.
- Supplementary Figure 6. Vaginal epithelial cells stained using a modified Harris-Schorr technique.
- Supplementary Figure 7. Composite profile demonstrating fluctuations in EI over the course of the ovarian cycle

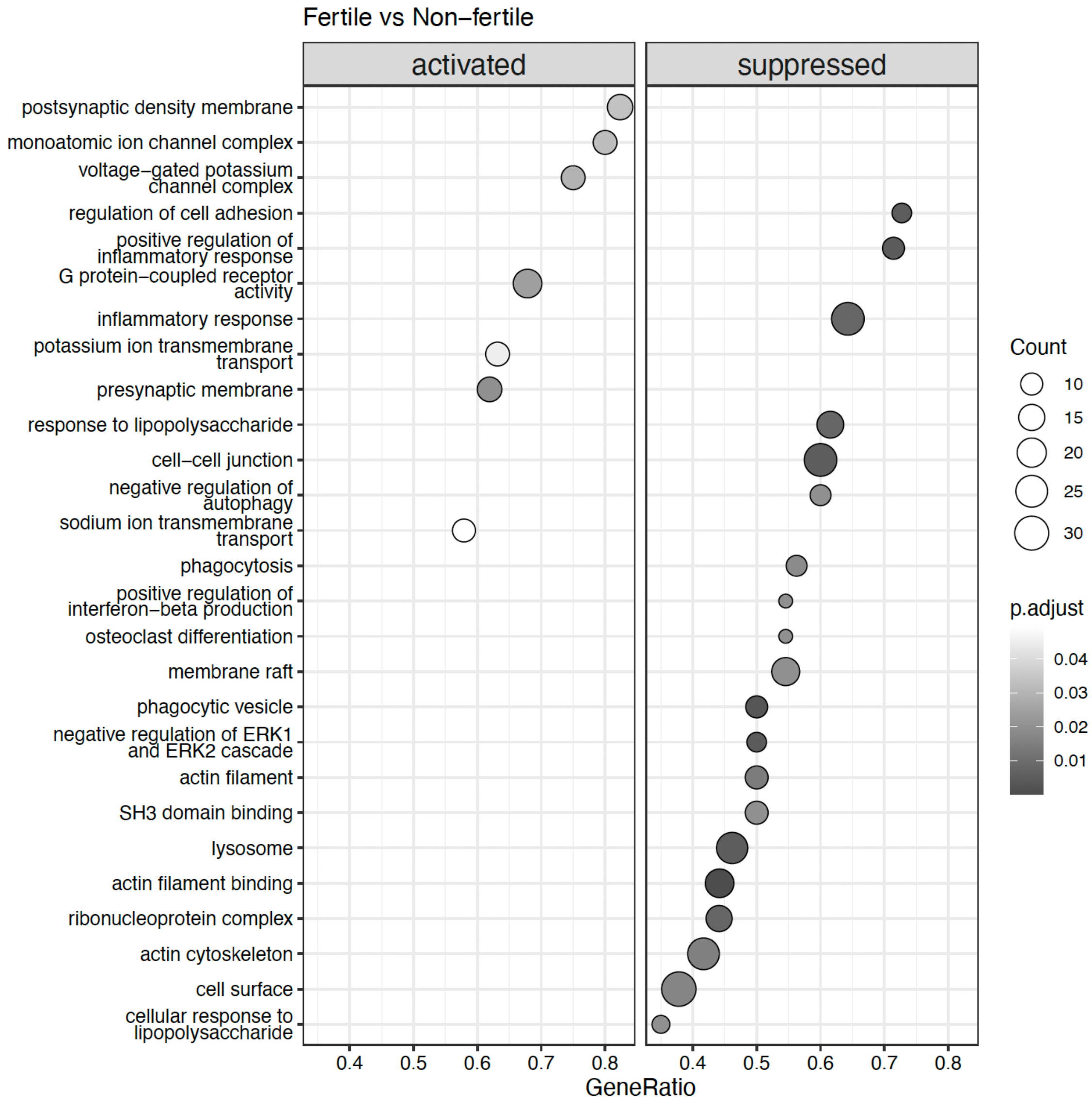


**Fig. S1. GSEA pathways enriched for differential expression in the fertile phase.** The "activated" panel represents pathways enriched for genes with heightened expression in the fertile compared to non-fertile phase and the "suppressed" panel represents pathways enriched for genes with lower expression in the fertile compared to non-fertile phase. Dot size represents the number of genes represented within that pathway and the darker colors represent lower p-values.

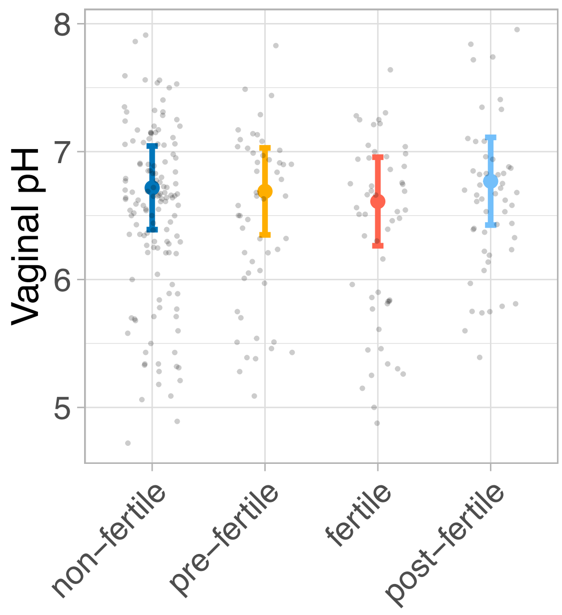


**Figure S2. Vaginal pH does not vary significantly between cycle phases.** Error bars represent model predictions +/- one standard error and points represent the raw data.

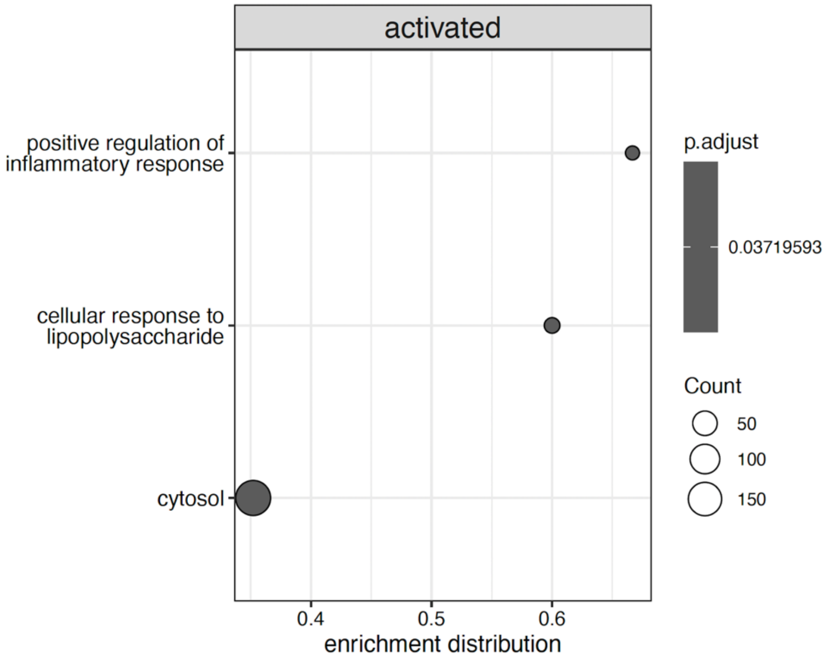


**Fig. S3. GSEA pathways enriched for differential expression post-copulation.** Three gene set pathways are enriched for genes with heightened expression in post-copulatory versus non-copulatory contexts. Dot size represents the number of genes represented within that pathway and the darker colors represent lower p-values.

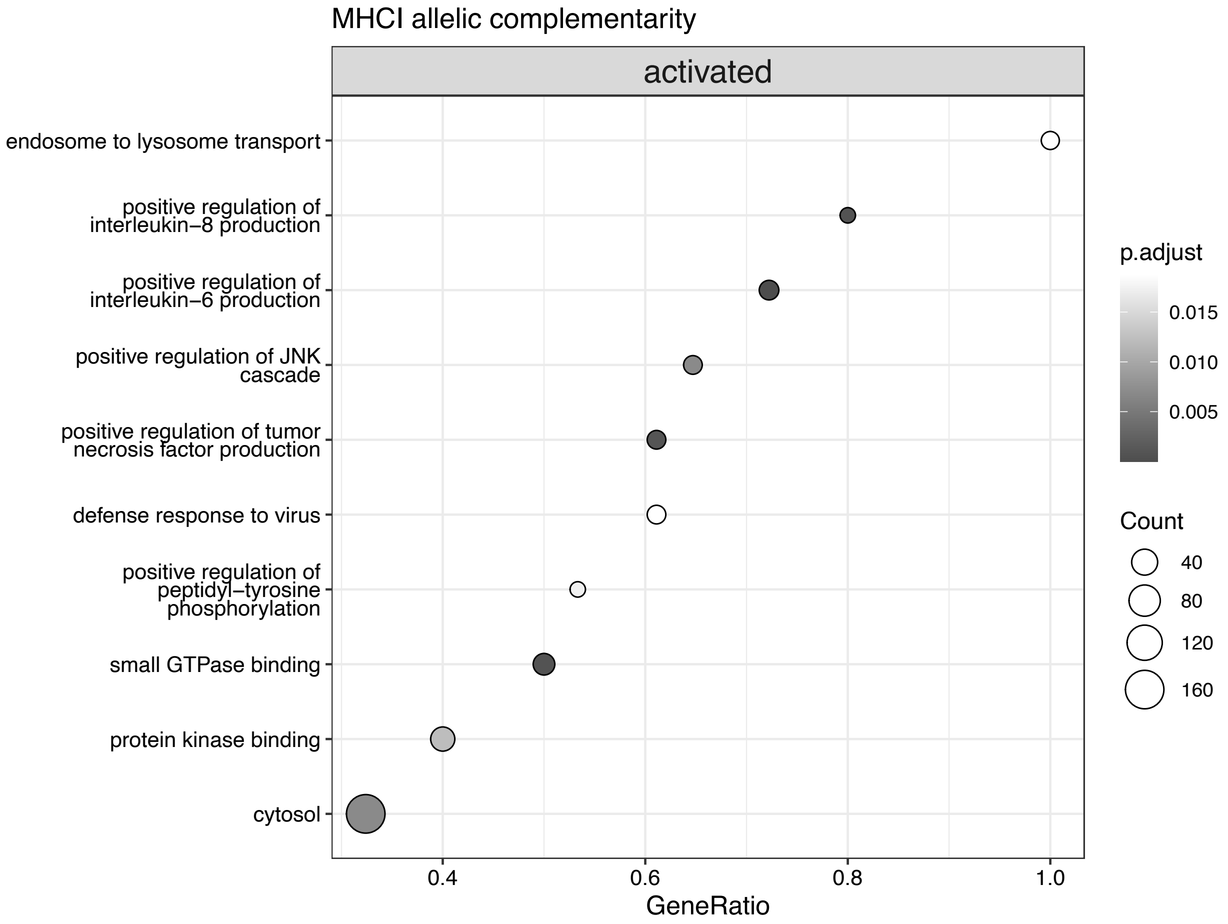


**Fig. S4. GSEA pathways enriched for differential expression in relation to MHC I allelic complementarity.** Pathways enriched for genes with heightened expression after mating with males with low complementarity. Dot size represents the number of genes represented within that pathway and the darker colors represent lower p-values.

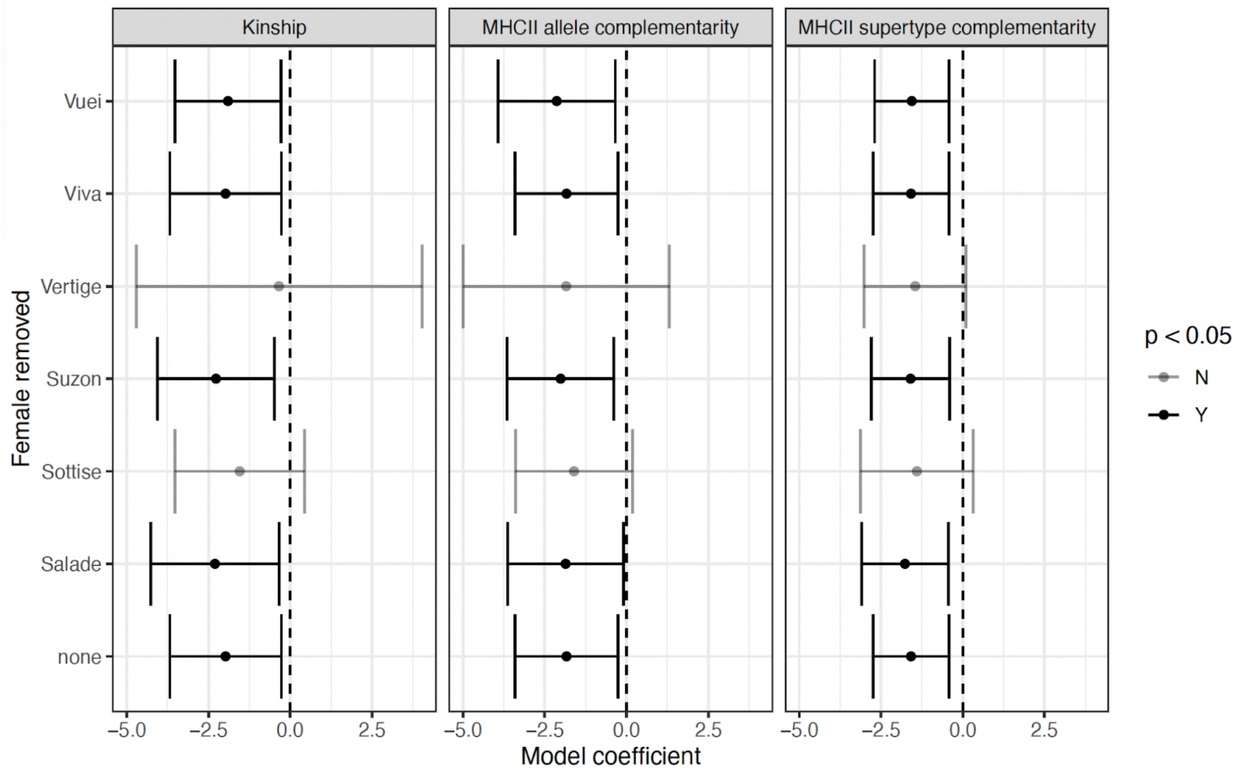


**Fig S5. Leave-one-out sensitivity analysis of vaginal pH model estimates.** Shown are estimated interaction effects between postcopulatory status and male genotype on vaginal pH across leave-one-out iterations. Points represent the estimated interaction term (postcopulatory status × male genotype), and error bars denote 95% confidence intervals. The genotype included in each model is indicated in the facet title.

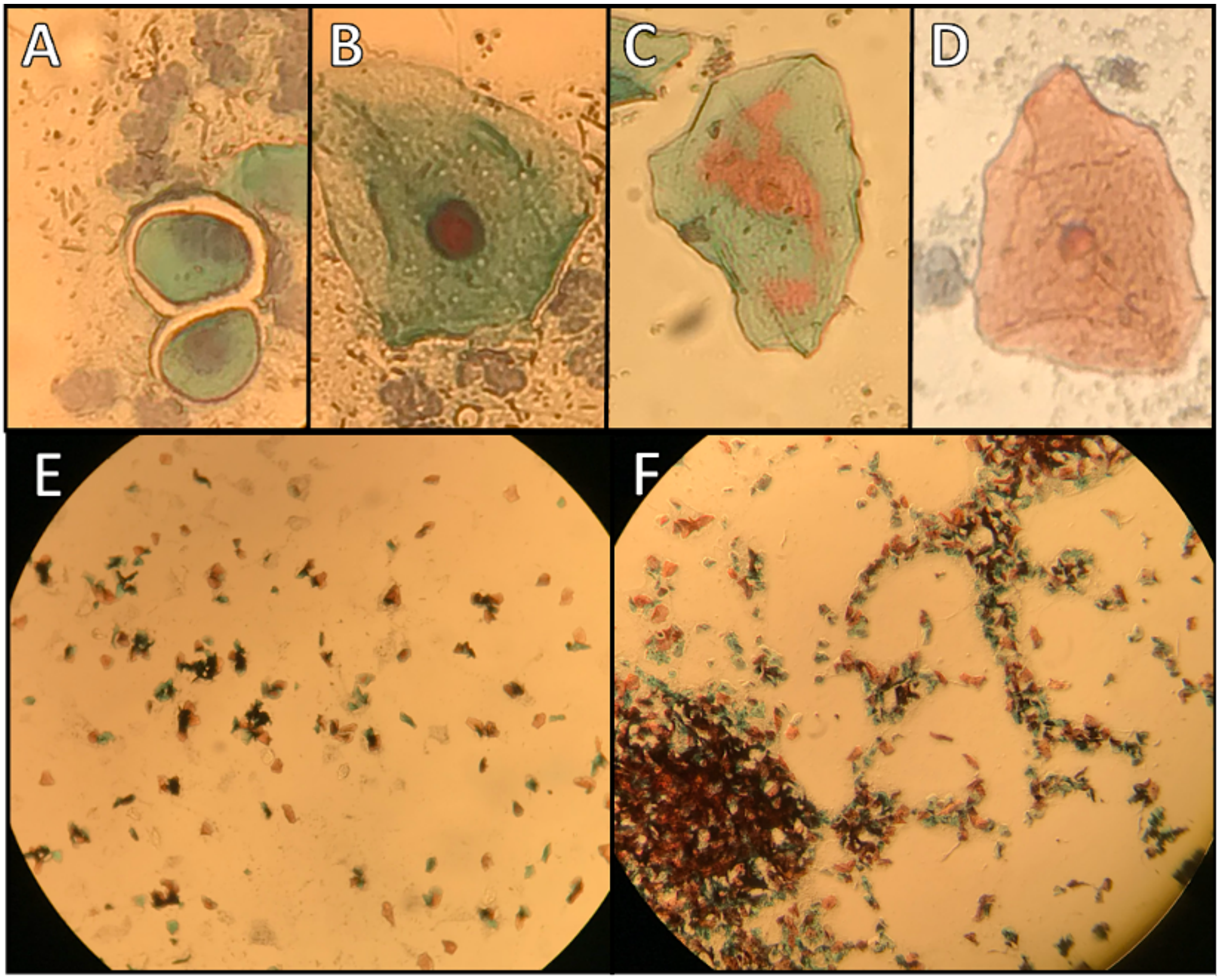


**Fig S6. Vaginal epithelial cells stained using a modified Harris-Schorr technique.** Epithelial cell types include basal cells (A), intermediate cells (B), polychromatophilic superficial cells (C), and eosinophilic superficial cells (D). The preovulatory phase (E) exhibits a gradual increase in the proportion of red to blue staining cells, and the postovulatory phase (F) is characterized by an increase in cellular clumping, mucus, and WBCs.

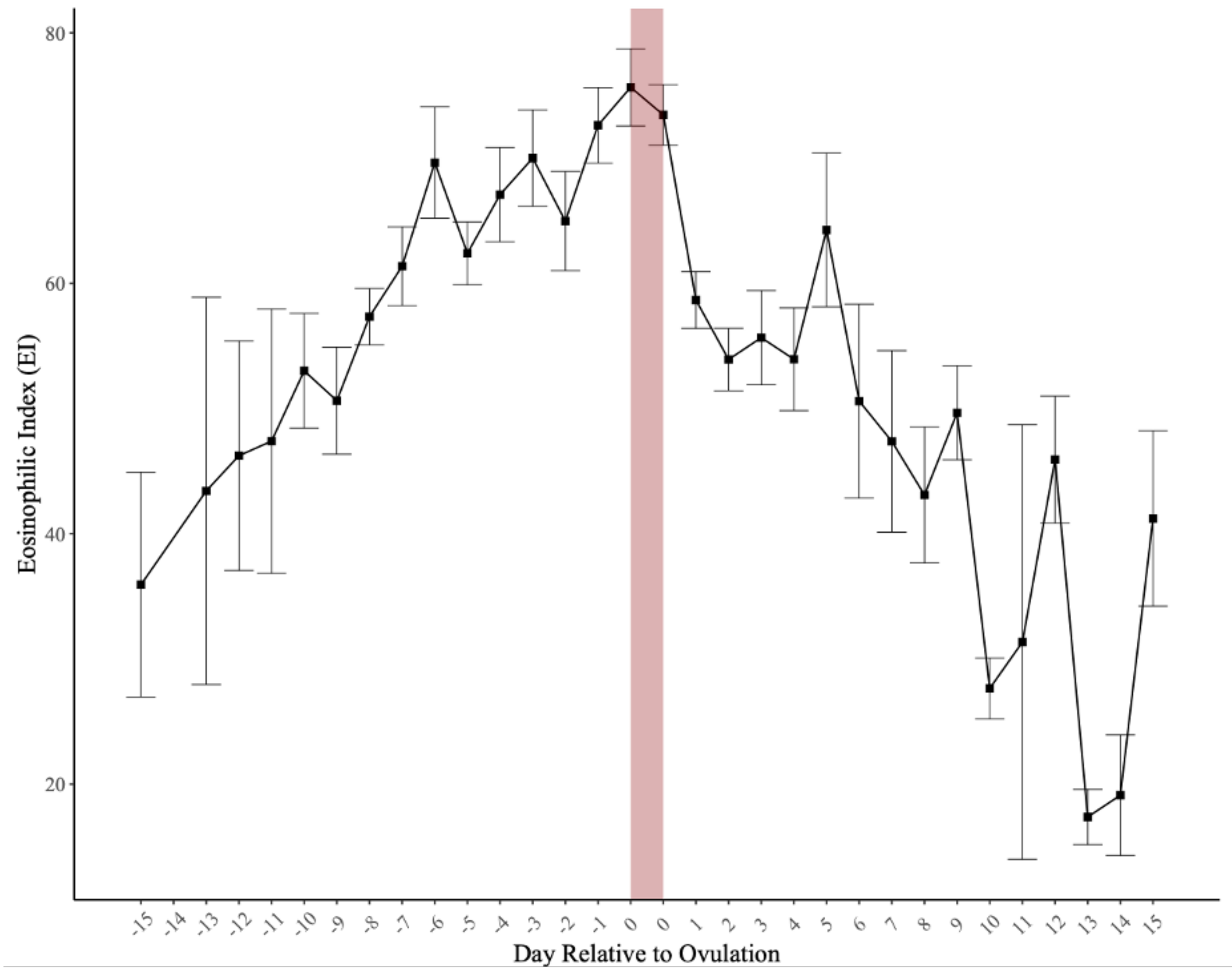


**Fig S7. Composite profile demonstrating fluctuations in EI over the course of the ovarian cycle (n= 22 cycles).** Points represent mean EI values on each day in relation to ovulation, and error bars represent the standard error of the mean. The two-day ovulation window is designated by consecutive 0's on the x axis and is shaded in red. The days leading up to ovulation are designated by negative numbers and the days following ovulation designated by positive numbers.
